# Fecal virome transplantation is sufficient to alter fecal microbiota and drive lean and obese body phenotypes in mice

**DOI:** 10.1101/2023.02.03.527064

**Authors:** Joshua M. Borin, Roland Liu, Yanhan Wang, Tsung-Chin Wu, Jessica Chopyk, Lina Huang, Peiting Kuo, Chandrabali Ghose, Justin R. Meyer, Xin M. Tu, Bernd Schnabl, David T. Pride

## Abstract

**Background:** The gastrointestinal microbiome plays a significant role in numerous host processes and has an especially large impact on modulating the host metabolism. Prior studies have shown that when mice receive fecal transplants from obese donors that were fed high-fat diets (HFD) (even when recipient mice are fed normal diets after transplantation), they develop obese phenotypes. These studies demonstrate the prominent role that the gut microbiota play in determining lean and obese phenotypes. While much of the credit has been given to gut bacteria, studies have not measured the impact of gut viruses on these phenotypes. To address this shortcoming, we gavaged mice with viromes isolated from donors fed HFD or normal chow. By characterizing the mice’s gut bacterial biota and weight-gain phenotypes over time, we demonstrate that viruses can shape the gut bacterial community and affect weight gain or loss.

**Results:** We gavaged mice longitudinally over 4 weeks while measuring their body weights and collecting fecal samples for 16S rRNA amplicon sequencing. We evaluated mice that were fed normal chow or high-fat diets, and gavaged each group with either chow-derived fecal viromes, HFD-derived fecal viromes, or phosphate buffered saline controls. We found a significant effect of gavage type, where mice fed chow but gavaged with HFD-derived viromes gained significantly more weight than their counterparts receiving chow-derived viromes. The converse was also true: mice fed HFD but gavaged with chow-derived viromes gained significantly less weight than their counterparts receiving HFD-derived viromes. These results were replicated in two separate experiments and the phenotypic changes were accompanied by significant and identifiable differences in the fecal bacterial biota. Notably, there were differences in Lachnospirales and Clostridia in mice fed chow but gavaged with HFD-derived fecal viromes, and in Peptostreptococcales, Oscillospirales, and Lachnospirales in mice fed HFD but gavaged with chow-derived fecal viromes. Due to methodological limitations, we were unable to identify specific bacterial species or strains that were responsible for respective phenotypic changes.

**Conclusions:** This study confirms that virome-mediated perturbations can alter the fecal microbiome in an *in vivo* model and indicates that such perturbations are sufficient to drive lean and obese phenotypes in mice.

## Background

In recent decades, discoveries in microbiome research have revealed myriad paths by which gut microbiota affect and regulate their hosts. Studies have demonstrated how resident microbes support health [1], as well as cause disease [2], and have revealed many of the processes by which microbiome communities can affect host immunity [3,4], metabolism [5], and even cognition [6,7]. Altogether, contributions to this body of work have catapulted an appreciation of the important and multifarious roles that resident microbiota play in their host’s life.

One seminal study, published in 2006 by Turnbaugh et al. [8], provides an early and compelling demonstration of the far-reaching effects of the microbiome. Researchers transplanted the gut microbiome from obese and lean mice into germ-free mice fed a normal chow diet. They found that individual mice receiving “obese” microbiomes gained more body fat and harvested more energy from the same quantity of food than mice receiving “lean” microbiomes. These results provide a striking demonstration of how microbiomes can alter their host’s metabolism to such a degree that it shapes their host’s phenotype.

At the time that this study was published, microbiome research was predominantly focused on the bacterial members of the microbiome. This focus led to many valuable contributions to the field. However, the gut microbiome is a diverse and complex community that also includes archaea, fungi, and viruses [9]. Over the last two decades, advances in sequencing technology and bioinformatic analysis have revealed that viruses in the gut can be more abundant than bacteria and that the gut virome is predominantly comprised of bacterial viruses called bacteriophages (phages) [10-12]. These discoveries have led to a burgeoning field investigating the host-associated virome and its effects on the microbiome and host organism.

Just as pioneering works shed light on the importance and influence of the gut bacterial biota, many studies now indicate that the gut virome can similarly affect its host’s health, metabolism, and even cognition and memory [11,13,14]. Shifts in virome composition have been associated with various conditions, including stunted growth [15], cancer and diabetes [12], and high-fat and high-sugar diets [16-18]. Furthermore, the administration of phages as treatments have been shown to attenuate the severity of bacteria-caused diseases, such as *Clostridium difficile* infections [19], *Klebsiella pneumoniae*-associated inflammatory bowel disease [20], and alcoholic liver disease caused by cytolysin-positive *Enterococcus faecalis* [21]. Fecal virome transplants (FVT) and gavages of specific phages with known bacterial hosts have also been shown to cause shifts in the bacterial composition of the gut [22-25] and bacterial metabolism [26], which could cause downstream effects on the host organism.

These discoveries have broadened our perspectives to include the virome as an important factor that shapes community assembly and host health. They have also led many to revisit interpretations from past fecal microbiome transplant studies; because viruses are smaller than bacterial cells, fecal microbiome transplants in previous studies likely contained viruses, in addition to bacteria. It is possible that many effects that were previously attributed to bacteria in fecal microbiome transplants could also be due to transplantation of associated viruses.

In this study, we explore whether viruses alone can alter the composition of the microbiome and drive changes in host phenotypes. To investigate, we conducted two longitudinal studies where mice on either normal chow or high-fat diets received fecal virome transplants prepared from donor mice on chow or high-fat diets (or phosphate buffered saline controls). By sampling mouse feces and body weights over time, we reveal how virome transplantation alters the composition of the gut microbiome and drives lean and obese phenotypes in recipient mice.

## Methods

### Animal study design

To study the effect of fecal virome transplantation (FVT) on microbiome composition and host animal physiology, we used a total of 72 male C57BL/6 treatment mice (Charles River Laboratories; Wilmington, MA) in two independent experimental replicates (i.e., trials; 24 mice in trial 1 and 48 mice in trial 2). In each trial, 6-week-old mice were randomly assorted into cages (4 and 3 mice per cage in trials 1 and 2, respectively). Half of the cages were given a standard laboratory chow diet and half were given a high-fat diet (HFD) (BioServ, S3282, 0.0341% cholesterol, 60% fat calories). After 10 weeks on respective diets, we administered fecal virome transplants via oral gavage every weekday for 4 weeks. Within each diet, cages were randomly assigned one of three gavage treatments: Chow-derived virome, HFD-derived virome, or a phosphate-buffered saline control (PBS). Feces from each cage were collected at regular intervals and stored at −80°C for downstream microbiome analyses. After 4 weeks of gavage treatment, mice were sacrificed and the weight of the liver and various adipose tissues were measured. Mice had ad libitum access to water, were housed under SPF conditions and maintained on a 12 h artificial light/dark cycle, a temperature range of 68–72°F, and a humidity of 40–70% RH. All animal studies were reviewed and approved by the International Animal Care and Use Committee of UCSD (#S09042).

### Fecal virome transplant preparation

To prepare material for FVT gavage, we fed a separate “donor” group of male C57BL/6 (Charles River; male, age 6 weeks) mice either chow or HFD for 10 weeks prior to starting gavage treatments (4 mice per cage). Virome gavages were prepared from the feces of mice on each diet every weekday: 0.5 g of donor feces were homogenized in 50 mL of SM buffer (100 mM NaCl, 8 mM MgSO_4_, 50 mM Tris-Cl pH 7.5 dissolved in water) and then serially passed through 0.45 μm and 0.2 μm filters to remove cells and debris. We confirmed that bacteria were successfully removed by inoculating 100 μL of filtrate into 4 mL of Luria-Bertani broth and incubating tubes at 37°C overnight. No observable growth was found. After filtration, the viruses in each sample were concentrated using polyethylene glycol (PEG) precipitation. Samples were subdivided into 15 mL conical tubes. Sterile NaCl and PEG-6000 solutions were added, bringing samples to a final concentration of 0.5M NaCl and 10% PEG-6000. Tubes were mixed and then stored overnight at 4°C. The following day, tubes were centrifuged at 4600 × *g* for 30 minutes to pellet the viruses. Then the supernatant was decanted and the pellet was resuspended in PBS. PEG-precipitated viromes within each diet type were pooled and stored at 4°C for no longer than 1 week. Before oral administration of the gavage, we added a 0.2 μm filter-sterilized solution of NaHCO_3_ to a final concentration of 5% to mitigate degradation of viruses by gastric acid.

### Quantification of viruses in gavage samples

Epifluorescence microscopy was used to enumerate the number of virus-like particles (VLPs) in PEG-precipitated gavage preparations. Samples were fixed in 2% formaldehyde, diluted, and vacuum filtered onto a 0.02 μm Anodisc filter. Next, the viral nucleic acids were stained by immersing the Anodisc in a droplet of 25X SYBR green I. After washing the Anodisc in a droplet of molecular grade water, it was mounted onto a microscope slide with 30 μL of antifade solution (1% p-phenylenediamine in 1:1 glycerol PBS) and imaged at 1000× via fluorescence microscopy. VLPs were manually enumerated in 3–5 fields of view per slide and the concentration of VLPs in each sample was calculated by N_v_ = P_t_ / F_t_ * A_t_ / A_f_ / V_t_ where N_v_ is VLPs per mL, P_t_ is the number of VLPs counted, F_t_ is the number of fields of view counted, A_t_ is the total area of the Anodisc, A_f_ is the area of each field of view, and V_t_ is the volume of sample that was filtered onto the Anodisc [27].

### 16S rRNA gene amplicon sequencing

To conduct 16S amplicon sequencing on feces collected during the study, DNA was first extracted using the Qiagen DNeasy Powersoil kit (Qiagen; CA). We included negative controls to ensure that no samples were contaminated during the extraction process. From purified DNA, we amplified the V3-V4 hypervariable region of the 16S rRNA gene via PCR with Hifi Hotstart Readymix (Kapa Biosystems; Boston, MA), forward primer 5’-TCG TCG GCA GCG TCA GAT GTG TAT AAG AGA CAG CCT ACG GGN GGC WGC AG-3’, and reverse primer 5’-GTC TCG TGG GCT CGG AGA TGT GTA TAA GAG ACA GGA CTA CHV GGG TAT CTA ATC C-3’ under the following cycle parameters: 95°C for 3 min, followed by 35 cycles of 95°C for 30 s, 55°C for 30 s, and 72°C for 30s, and a final elongation of 72°C for 5 min [28]. PCR reactions were cleaned using AMPure XP beads (Beckman-Coulter; Fullerton, CA) and then indexed using a Nextera XT DNA Library Preparation Kit (Illumina; San Diego, CA). Finally, samples were cleaned again with Ampure XP beads, visualized with a High Sensitivity DNA Kit on a Bioanalyzer (Agilent Technologies; Palo Alto, CA) and quantified using a dsDNA High Sensitivity Kit on a Qubit Fluorometer (Thermo Fisher Scientific; USA). Samples were pooled into equal molar proportions and sequenced on the Illumina MiSeq platform (Illumina; San Diego, CA).

### Analysis of 16S rRNA sequences

Reads were processed using Quantitative Insights Into Microbial Ecology 2 (QIIME2; version 2021.4) [29]. Quality filtering, dereplicating, and removal of chimeras was handled by the DADA2 plugin in QIIME2 [30]. Taxonomy classification was performed using the feature-classifier “classify-sklearn feature” in QIIME2, with a Naïve Bayes classifier trained on the SILVA database (version 138) [31]. The relative abundance of different bacterial classes was visualized using the qiime2R (v0.99) and ggplot2 (v3.3.5) packages in R (v4.1.1) [32-34]. Alpha diversity (Shannon index, Faith’s Phylogenetic Diversity, and Operational Taxonomic Unit) and beta diversity (Bray-Curtis) metrics were produced by the QIIME2 “core-metrics-phylogenetic” pipeline (sampling depth=8000). To further investigate the compositional differences between samples, we used the QIIME2 DEICODE plugin to compute Robust Aitchison distances and to create taxonomic biplot overlays.

### Virome shotgun sequencing

To sequence the fecal viromes of donor and recipient mice,viral DNA and RNA was extracted using an adapted “NetoVIR” protocol [35], adapted from [36]. First, 0.05 g of fecal material was homogenized in 500 μL of 0.5x PBS for 1 min, centrifuged at 17,000 x *g* for 3 min. The supernatant was then filtered through a 0.45 cellulose-acetate μm filter for 1 minute at 17,000 x *g*. Next, 130uL of filtered samples were nuclease treated with 2 μL of benzoase and 1 μL of micrococcal nuclease in 7 μL of 20x homemade buffer (1 M Tris pH 8, 100 mM CaCl_2_ 30 mM MgCl_2_). After gently pipetting to mix, samples were incubated at 37°C for 2 h and then the reaction was stopped by adding 7 μL of 10 nM EDTA. Finally, viral nucleic acids (both RNA and DNA) were extracted with the QIAamp Viral RNA Minikit (Qiagen; CA).

Nucleic acids were amplified by using the Complete Whole Transcriptome Amplification Kit Protocol, which converts viral RNA into cDNA prior to amplification of both the resulting cDNA and extracted viral DNA (WTA2, Sigma-Aldrich; USA). In order to capture rare viral nucleic acids, we increased the number of amplification cycles to 22. After amplification, we measured the DNA concentration using the Qubit dsDNA HS Assay Kit (Thermo Fisher Scientific; USA). Library preparation was performed using the Nextera XT DNA Library Preparation Kit (Illumina; San Diego, CA) with 1.2 ng/μL of input material. Tagmentation time was shortened to 4 min and PCR extension time increased to 45 s in order to favor larger DNA fragments. Finally, samples were purified using 0.6X AMPure XP beads (Beckman-Coulter; Fullerton, CA) and library size was checked using a Bioanalyzer (Agilent Technologies; Palo Alto, CA). Libraries were pooled into a 4 nM solution and sequenced on the Illumina MiSeq platform (Illumina; San Diego; CA) with a target sequencing yield of 500MB-1GB per virome.

### Analysis of virome sequences

Virome sequencing reads were analyzed using a modified version of the protocol described by Santiago-Rodriguez et al. (2015) [37]. First, reads were trimmed based on size (>100bp) and quality (>Q30) using CLC Genomics Workbench (Qiagen; CA). Reads were then mapped to common bacterial contaminants using CLC Genomics Workbench; these reads were removed before assembly. Reads were also mapped to the mouse BALB/c genome. As less than 0.1% of reads mapped to the mouse genome, these reads were not removed in case they represented endogenous murine viral sequences. We then assembled reads using CLC Genomics Workbench De Novo Assembly tool with 98% identity and a minimum 50% read overlap. Consensus sequences were constructed according to the majority rule. Contigs of short lengths (<500bp), containing ambiguous characters, and mapping to mouse mitochondrial DNA were removed. The remaining virome contigs were annotated using tBLASTx against NCBI All Viruses database with an E value cutoff of 10^−10^. Finally, the tBLASTx results were parsed via Ion Assist for classification of viral families.

### Statistics

Analyses on mouse body weights were modeled both cross-sectionally and longitudinally by generalized estimating equations [38]. For beta diversity, PERMANOVA (999 permutations) were conducted to test group differences based on the Bray-Curtis and Robust Aitchison diversity distance metrics, respectively [39]. Because alpha diversity values had nonlinear changes over time, we used a spline regression to capture such nonlinear changes, with three knots at days 4, 9, and 16 [40]. We then conducted Wald tests to compare Shannon index values between treatments on a given day, as well as to compare how the slope of alpha diversity in each treatment changed with respect to each knot. For multiple comparisons, p-values were corrected using the Holm method [41]. All statistical analyses were carried out in R [34].

## Results

### Experimental design and effect of FVT on mouse weight

To investigate how viruses in the gut influence host physiology and the gut microbiota, we administered fecal virome transplants (FVT) to mice in a 2-diet type × 3-gavage type experimental design (Fig. 1). Two independent experimental trials were conducted, consisting of 72 mice in total. First, recipient mice were acclimated to either normal chow or high fat diet (HFD) for 10 weeks. Then, we administered FVT (or PBS controls) via oral gavage every weekday for 4 weeks. Viromes for FVT were prepared from the feces of a separate group of donor mice that were fed chow or HFD for 13 weeks. To prepare viromes for gavage, feces from each diet type were suspended in buffer, homogenized, and filtered to remove cells. Finally, viruses were concentrated via PEG precipitation and resuspended in PBS. Each weekday, mice received an oral gavage containing ~5 × 10^9^ virus-like particles (VLPs). VLPs were enumerated via epifluorescence microscopy (Fig. S1). We found no significant difference in VLP concentration between HFD-derived and chow-derived preparations (1.73 × 10^10^ VLP/mL ±7.43 × 10^9^ SD and 1.02 × 10^10^ VLP/mL ±9.47 × 10^9^ SD, respectively).

**Figure 1.**
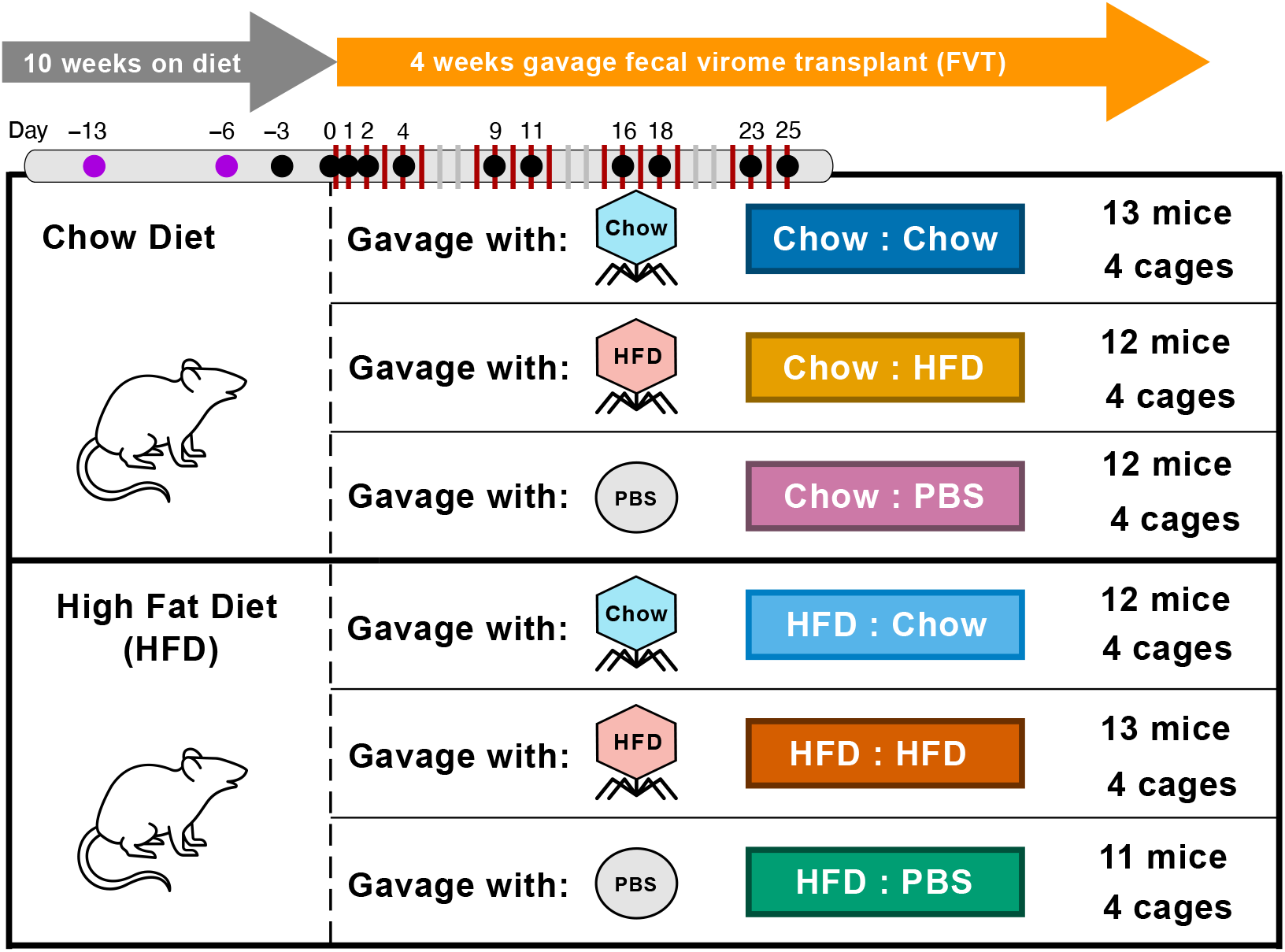
Study design. Recipient mice were fed chow or high-fat diet (HFD) for 14 weeks. After the first 10 weeks, fecal virome transplants (FVT) were administered via oral gavage every weekday for 4 weeks (red vertical bars) and feces were collected at regular intervals (dots). FVT were prepared in parallel from a separate group of donor mice on chow or high-fat diet. Gavage treatments consisted of chow-derived and HFD-derived viromes, as well as PBS controls. The study was conducted in 2 experimental replicates (i.e., trials). Trial 1 consisted of 4 mice per cage and 1 cage per treatment. Trial 2 consisted of 3 mice per cage and 3 cages per treatment. In Trial 2, we collected feces from 2 additional pre-gavage timepoints (purple dots). Five mice were omitted from the study due to death (one in Trial 1 HFD : PBS, one in Trial 2 HFD : PBS) or aggression toward other mice (one in each Trial 2 HFD : Chow, Chow : PBS, and Chow : HFD).

To explore how FVT affects host physiology, we recorded the body weights of individual mice two times per week after starting gavage treatments. As expected, mice fed a high-fat diet gained more weight than mice fed chow diets by the end of the experiment (Fig. 2A, p<2.2e-16). Surprisingly, we found that FVT significantly affected the amount of weight mice gained or lost during the 4-week experiment (Fig. 2A, p=0.0023). Mice receiving chow-derived viromes (prepared from the feces of donor mice on a chow diet) gained less weight than mice receiving HFD-derived viromes, regardless of the recipient mouse’s diet.

**Figure 2.**
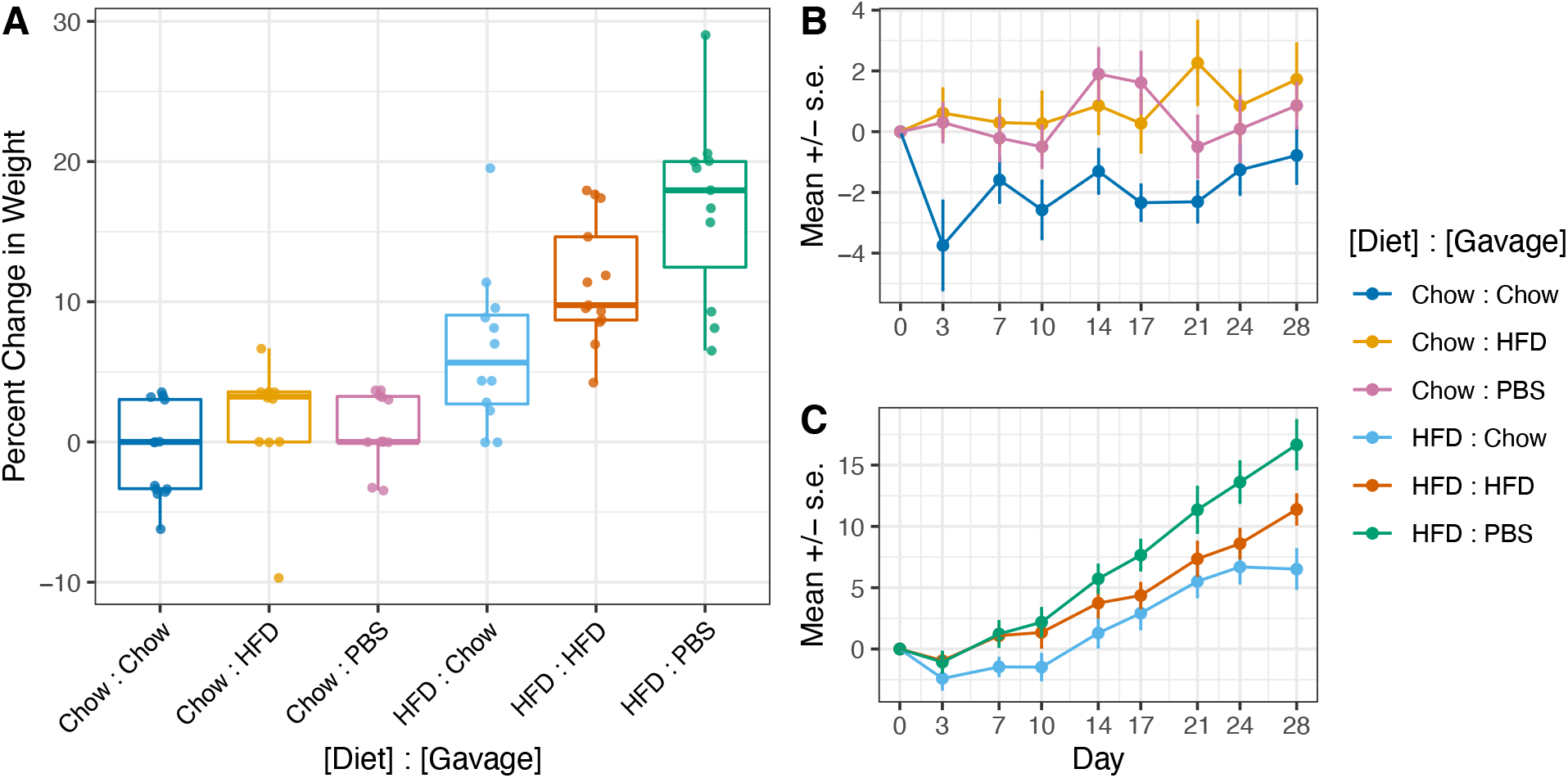
Change in weight of individual mice in response to diet and FVT during 4 weeks of gavage. Percent change in weight is shown from initial to final weight of individual mice (A), as well as the percent change in weight over time for mice fed chow (B) or HFD (C). For mouse weights after 28 d (A), Diet and Gavage effects are significant (p=2.2e-12, p=0.0023, respectively) and effects of Trial, Cage, and Diet-by-Gavage interaction are not. For longitudinal analysis (B and C), Diet, Gavage, and Diet-by-Time interaction are significant (p=0.028, p=0.031, p<2e-16, respectively) but Gavage-by-Time interaction is not. Data from PBS controls were omitted from statistical models.

Longitudinal analyses on how body weights changed over the 4-week experiment provided further insight into how FVT affects mouse physiology (Fig. 2B, Fig. 2C). Regardless of diet type, mice receiving chow-derived viromes lost more weight than mice on the same diet receiving HFD-derived viromes. By day 3, mice on a chow diet and receiving chow-derived viromes (blue, Fig. 2B) lost an average of 3.7% body weight (±1.46 SE) and mice on a high-fat diet receiving chow-derived viromes (light blue, Fig. 2C) lost an average of 2.4% body weight (±0.87 SE). However, after this initial perturbation, the change in body weight of mice receiving chow-derived viruses was similar to mice receiving other gavage treatments within the same diet type. These results are supported by longitudinal statistical analyses indicating that the initial effect of gavage is statistically significant (Gavage, p=0.031) but that this effect does not change over time (Gavage-by-Time, p=0.844), despite continued administration of FVT for 3 additional weeks.

At the end of the gavage period, mice were sacrificed to determine whether treatments affected the liver and various adipose tissues. We fit a GEE model to identify the relationship between tissue weight proportions and different combinations of diet type and gavage. As expected, we found a significant effect of diet (p<2.2e-16) for all of the tissues we measured (liver, epidydimal white adipose tissue, subcutaneous white adipose tissue, and brown adipose tissue), however, we did not detect a significant effect of gavage on any of these tissue weights (Fig. S2).

### Effect of FVT on gut bacterial community composition

To investigate how FVT affected the gut bacterial community of mice, we sequenced 16S rRNA amplicons from the feces of FVT recipient mice throughout the 4-week study. In total, 339 fecal samples were included in our analysis (16 donors, 86 from trial 1, and 237 from trial 2) with a total of 29,530,059 sequencing reads and an average of 86,628 reads per sample (± 22,078 SD). Because of unequal sequencing depth, samples were rarefied to a minimum sampling depth of 8,000 sequences per sample. This allowed us to capture the overall diversity within our samples without omitting too many from the study (Fig. S3).

We first investigated whether there were compositional differences in gut bacterial communities between treatments by computing Bray-Curtis dissimilarities. Data were visualized via Principal Coordinate Analysis as a complete dataset (Fig. 3A), as well as by diet and trial (Fig. 3B-E). Statistical analyses were conducted on the complete dataset using PERMANOVA (Fig. 3F). We found that there was a clear distinction between the bacterial communities of mice on different diets (R^2^=0.331, p=0.001), as well as between mice that were conducted in the first and second trials (R^2^=0.168, p=0.001) (Fig. 3A). We also found a significant effect of FVT on composition of the bacterial community (R^2^=0.018, p=0.001). Importantly, this effect did not persist over time (p>0.05), mirroring our analyses on mouse body weights. Detection of this effect was not driven by a single gavage type, as the gavage effect was significant (p<0.001) and explained a similar amount of the variance in all pairwise gavage comparisons (R^2^≈1.5%).

**Figure 3.**
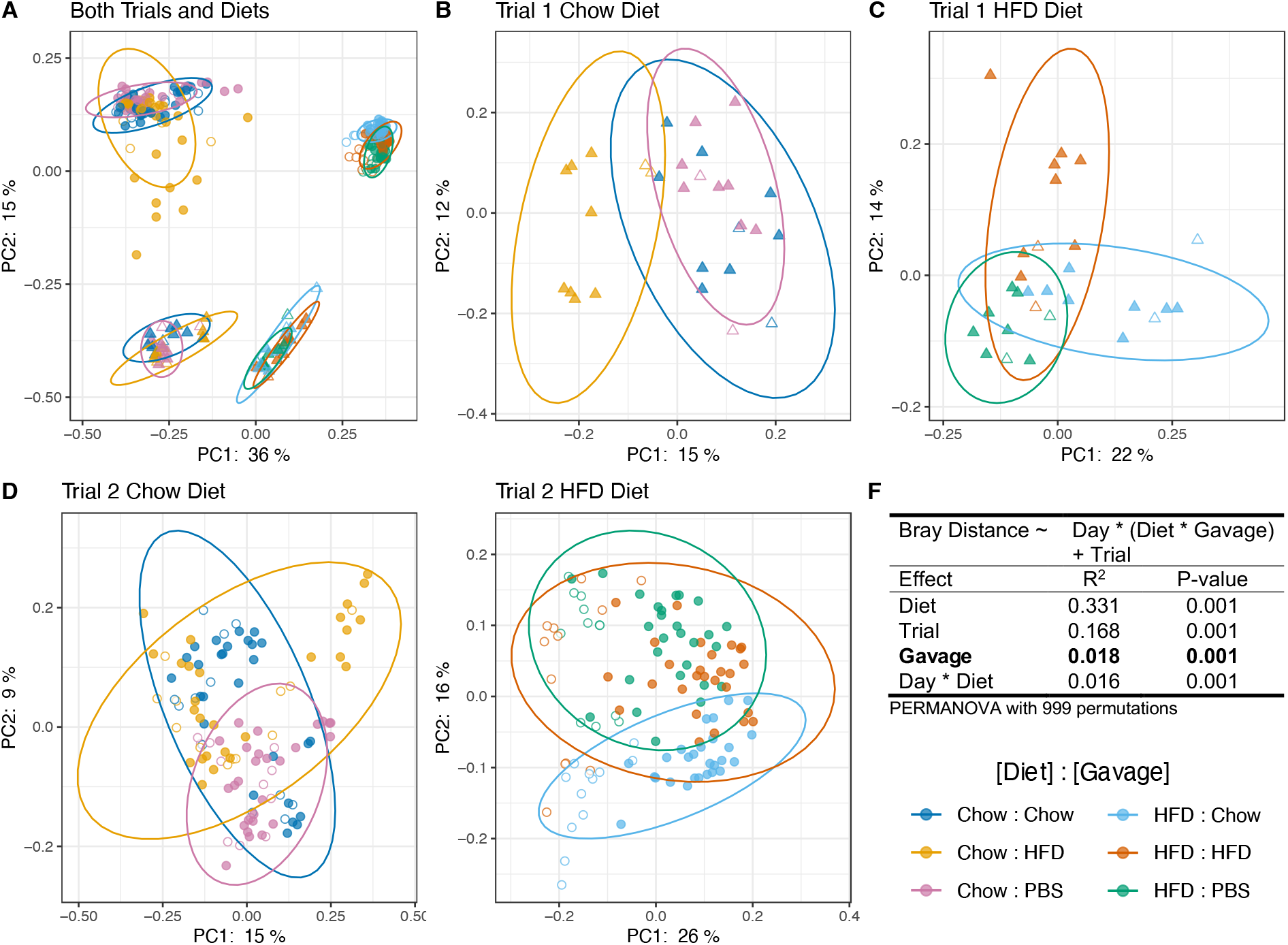
Bray-Curtis beta diversity of fecal microbiomes of FVT recipient mice visualized by Principal Coordinate Analysis (PCoA). Panel A: All recipient mice. Panels B and C: Trial 1 recipient mice fed chow (B) or HFD (C). Panels E and F: Trial 2 recipient mice fed chow (D) or HFD (E). Diet, Trial, Day, Gavage, and Day*Diet effects are significant when including all recipient mice (PERMANOVA with 999 permutations, R^2^ and p-values reported in Panel F). The same effects are significant for PERMANOVA conducted on all pairwise combinations of gavage treatments (HFD vs Chow, HFD vs PBS, Chow vs PBS). Unfilled points indicate pre-gavage samples collected before day 0. Statistical tests were conducted on data from day 0–25.

Next, we investigated whether FVT affected overall bacterial diversity in the gut by computing Shannon Index values across time (Fig. 4). We found that the microbiomes of mice consuming a high-fat diet were less diverse than those on a chow diet, consistent with previous studies [18]. We also discovered a significant effect of virome transplantation on alpha diversity; For trial 2 mice receiving a high-fat diet and HFD-derived viruses (orange line, Fig. 4B), Shannon Index values were significantly lower from days 9–18 compared to mice on the same diet receiving chow-derived viruses or PBS controls (p<0.05, Wald test with p-values adjusted for multiple comparisons). Interestingly, it took several days for HFD-derived viruses to induce detectable changes in alpha diversity and this effect eventually receded ~23 d after starting gavage treatments. We see a similar result in mice from trial 1 (orange line, Fig. 4A), however we lacked treatment replicates to conduct statistical analyses.

**Figure 4.**
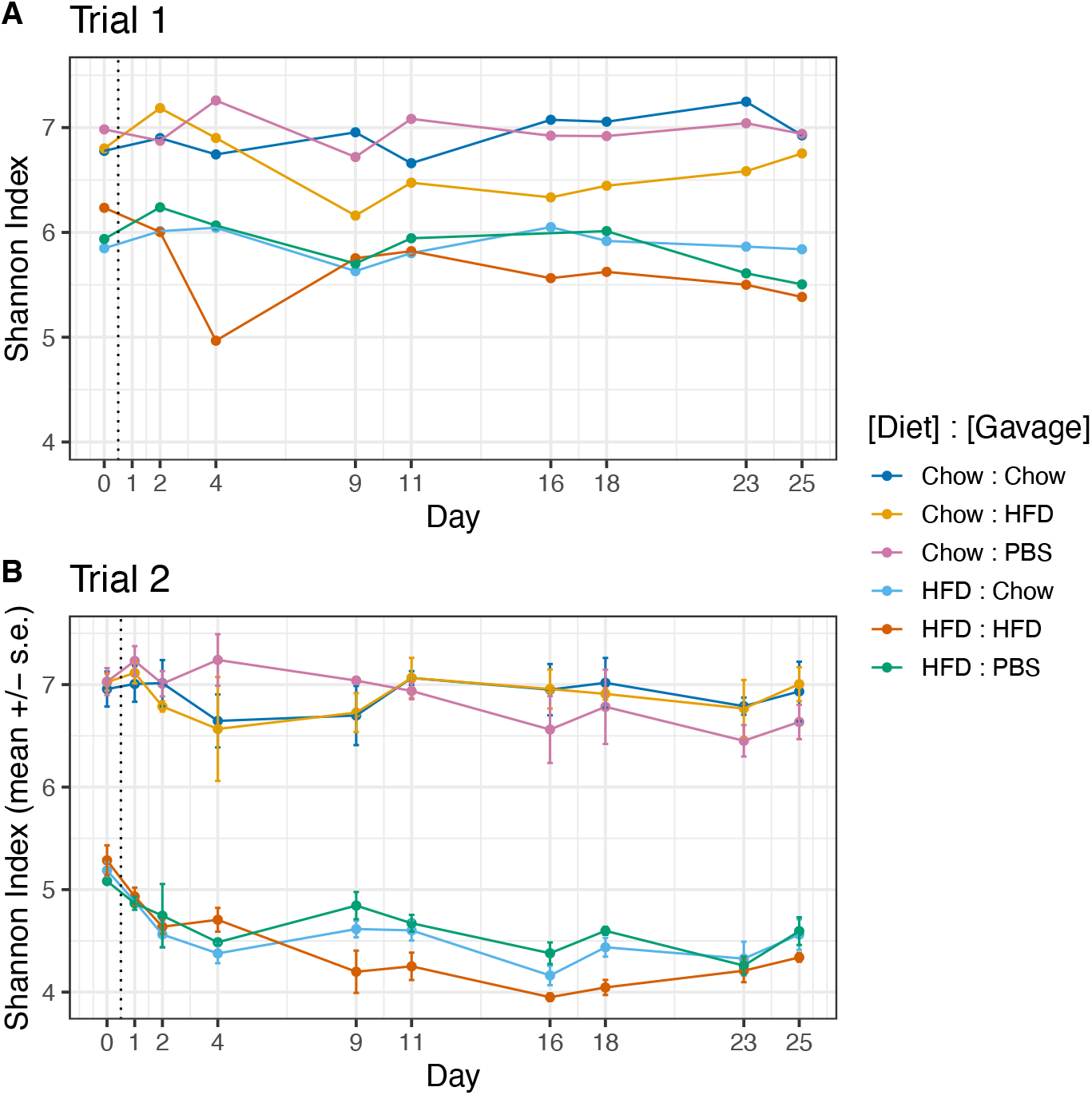
Shannon Index alpha diversity of fecal microbiomes during 4 weeks of FVT gavage from trial 1 (Panel A) and trial 2 (Panel B). Trial 1 (A) consisted of a single replicate per treatment, precluding statistical analyses. For trial 2, mice on HFD and receiving HFD-derived viromes (orange) had significantly lower alpha diversity between days 9–18 than mice on the same diet receiving chow-derived viromes (light blue) or PBS controls (green) (p<0.05, spline regression and Wald test with p-values corrected for multiple comparisons). Overall, mice on a high fat diet had lower alpha diversity that mice on a chow diet.

For mice on either diet receiving chow-derived viromes, we did not find an effect of gavage on alpha diversity (light and dark blue lines, Fig. 4). This was surprising, given that these mice showed the greatest reduction in body weight (light and dark blue, Fig. 2). Therefore, we did not find any correlation between how FVT affects alpha diversity and how it affects body weight.

To identify which bacterial taxa were affected by FVT, we first visualized the relative abundances of bacterial classes in recipient mice throughout the gavage period. In trial 2 (Fig. 4), trial 1 (Fig. S4), and for trial 2 donor mice (Fig. S5), gut microbiomes were predominantly comprised of Clostridia (blue) and Verrucomicrobiae (orange), and many samples contained visible fractions of Alphaproteobacteria (red), Coriobacteria (light blue), and Saccharimonadia (yellow). Mice in trial 2 on a high-fat diet (Fig. 4, bottom row) show reproducible increases in Verrucomicrobiae and Alphaproteobacteria coinciding with the beginning of gavage treatment. However, these increases also appeared in the PBS treatments, suggesting that this could be caused by the gavage procedure and not by transplanted viruses. No reproducible increase or decrease in bacteria was associated with a specific gavage type at the “class” level. Therefore, we hypothesized that FVT may have affected the microbiome and mouse body weight via slight alterations in the abundance of multiple taxonomic groups.

To explore whether FVT exerted an effect by slightly altering multiple bacterial taxa, we adopted a second beta diversity metric using Aitchison principal component analysis (PCA). Importantly, Aitchison distances are robust to high levels of sparsity and can be used to reveal inter-community differences when visualized as compositional biplots [42]. Because community composition was distinct between diet types and experimental trials (see Fig. 3A, and Fig. 3F), we separated the data by diet and trial and then computed Aitchison distances and conducted statistical analyses.

We find that, when considering mice within a given diet and trial, the effect of gavage explains an average of 36.5% (range 20–44%) of the variance between samples. In addition, Aitchison PCA reveals the 8 taxonomic groups that explain the majority of dispersion between samples in the biplot (Fig. 5). Interestingly, all of the bacterial orders identified belong to the class Clostridia (Clostridia UCG0-14, Lachnospirales, Oscillospirales, Peptostreptococcales, Ruminococcaceae), except for Rhodospirillales which belongs to class Alphaproteobacteria. Overall, these results suggest that phages in the FVT primarily affected the most abundant bacterial class (Clostridia, see Fig. 4). However, they also reveal that members of a given bacterial order were not all affected in the same way. For example, in trial 1 chow diet mice, some groups of Lachnospirales were more abundant under HFD FVT and other groups of Lachnospirales were more abundant under chow FVT (Fig. 5A). Altogether these results suggest that FVT induced changes, primarily to the most abundant bacteria in the mouse gut, and that these changes can ultimately alter the physiology of the host mouse.

**Figure 5.**
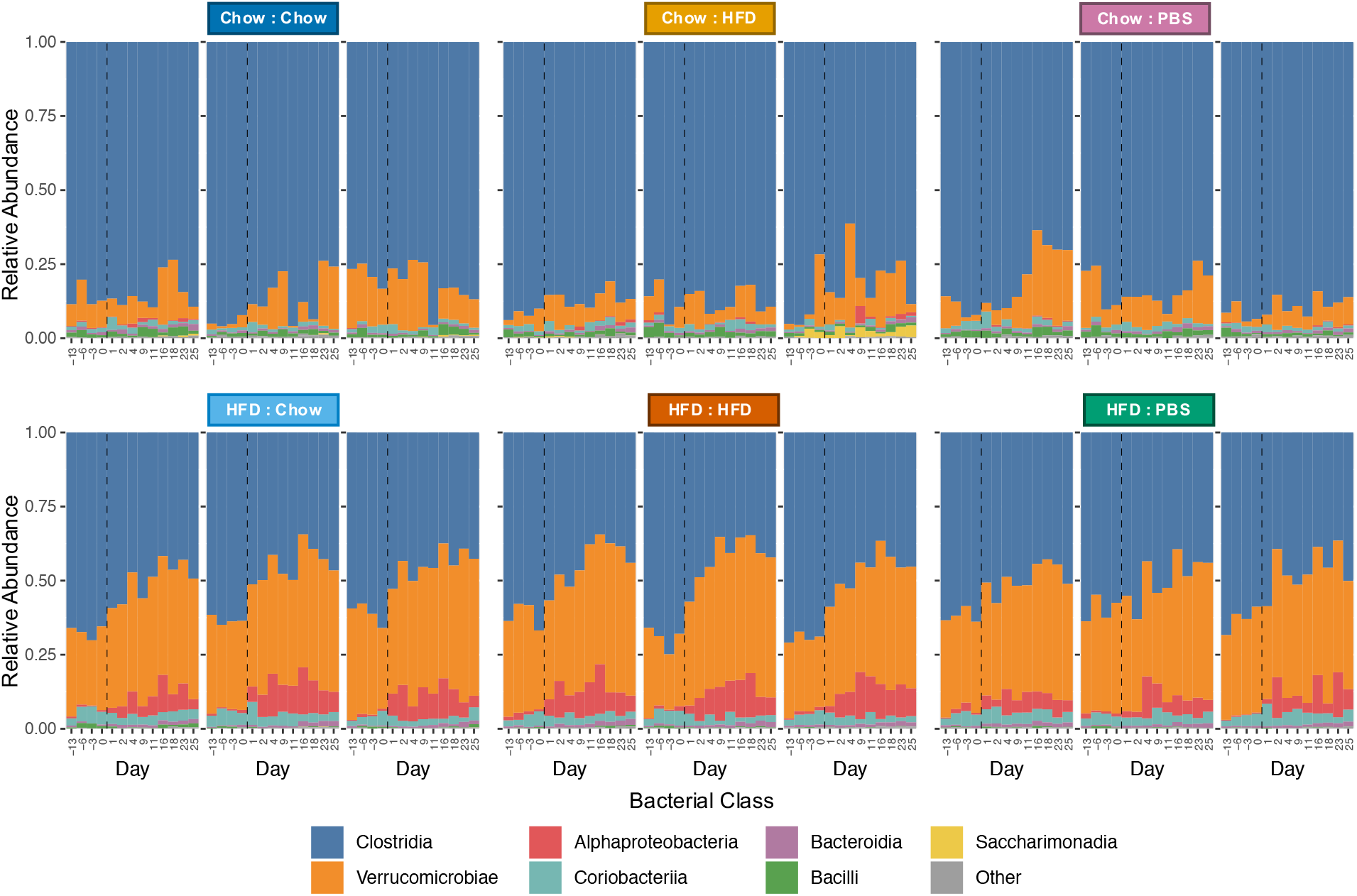
Taxonomy stacked bar plots of bacterial classes throughout the 4-week experiment. Each panel represents a cage of 3 mice. Replicate cages (n=3) are grouped and labeled by treatment (Diet: Gavage). Recipient mice received FVT gavage prepared from donor mice on chow diet or HFD, or a PBS control. The dashed line indicates start of gavage treatment.

**Figure 6.**
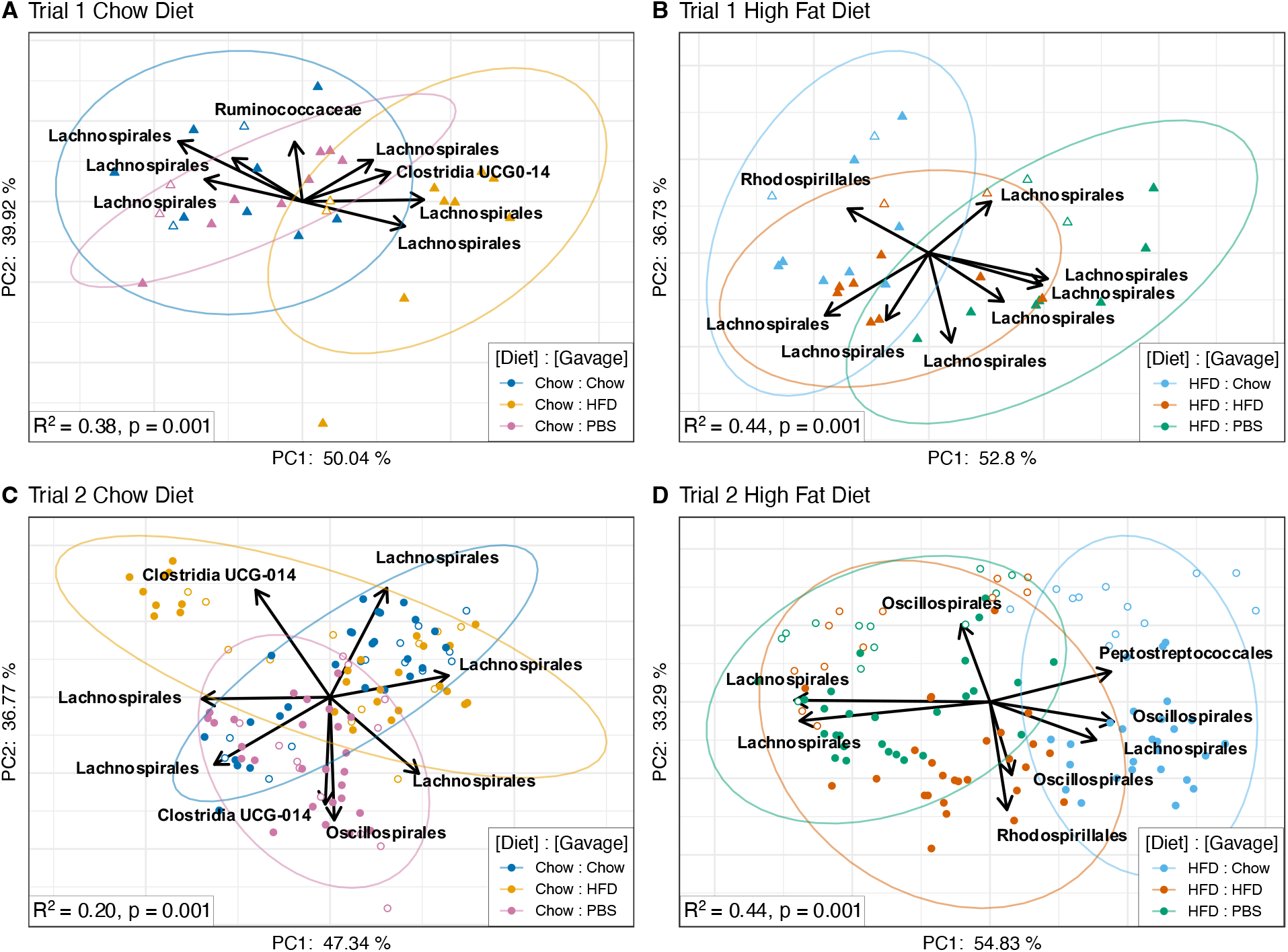
Aitchison compositional biplots of recipient mice visualized by principal component analysis (PCA). Panels A and B: Recipient mice from trial 1 fed chow (A) or HFD (B). Panels C and D: Recipient mice from trial 2 fed chow (C) or HFD (D). Color denotes treatment group (see Fig. 1). Arrows denote important taxa in relation to sample clusters. Significance was determined by PERMANOVA with 999 permutations (Aitchison distances ~ Day * Gavage); The R^2^ and p-value for the effect of gavage are indicated in respective panels.

## Discussion

The discovery that fecal microbiome transplants can shape host metabolism and impact lean and obese phenotypes in mice was a seminal breakthrough in microbiome research [8]. At the time of its publication, little consideration was given to the host virome and the fact that fecal microbiome transplants contain myriad viruses. Since then, several groups have shown that viral communities are altered in certain disease conditions such as periodontal disease [43] and inflammatory bowel disease [44]. Yet, it remains unclear whether differences in viral communities were a reflection of the changes in bacterial communities or whether viruses were *responsible* for the observed changes. To address this issue, we set out to determine whether viruses were sufficient to alter the microbiome and potentially affect lean and obese phenotypes in mice.

Using previously described techniques [35], we purified and then transplanted fecal viromes into mice on different diet types via oral gavage. In order to determine whether viruses directly affected mouse phenotypes, we recorded mouse body weights throughout 4 weeks of study. We observed the same phenomenon in two separate trials: mice on chow diets gavaged with HFD-derived viruses gained significantly more weight, and mice on high-fat diets gavaged with chow-derived viruses gained significantly less weight. These data indicate that virome gavages, which are devoid of bacteria, are sufficient to alter host metabolic phenotypes.

To explore how virome gavages altered the gut microbiota, we conducted 16S rRNA amplicon sequencing and analyses on fecal samples collected longitudinally throughout the study. Given that fecal virome gavages were largely comprised of phages, it is not surprising that transplanted viruses showed a capacity to affect the gut bacterial community. This study is not the first to demonstrate that the microbiota can be shaped by phages; Virus-mediated perturbations have been shown to impact oral, gut, and model-gut microbiota [22,25,35]. Through their effect on the microbiota, phage treatments have also been shown to affect aspects of host metabolism [22,24,26], and attenuate diseases such as *Clostridium difficile* colitis [19] and alcoholic liver disease [21]. The aforementioned studies used a diversity of virus perturbations, including single-phage treatments, synthetic assemblages of phages, and unaltered virus communities. Altogether, our work, as well as previous studies highlight the roles that viruses play in shaping bacterial communities and affecting disease and metabolism of the host organism.

While virome gavages were sufficient to alter the body weight of mice, it is unclear whether the extent of weight gain/loss in our study is similar to that which was observed by Turnbaugh et al. (2006) [8], whom used full fecal microbiome transplants. We believe that the degree of weight gain/loss in this study is likely different from those in Turnbaugh et al., however further study is needed to directly compare the results of both protocols. In mice, it is relatively straightforward to identify determinants of weight gain [45] and our results reveal a clear trend between fecal virome transplantation and mouse body weight. However, it is unclear whether virome transplants can affect body weight in humans and further study is necessary to corroborate such a suggestion.

In humans, it has been challenging to identify microbes that are reproducibly associated with weight-gain and weight-loss. In this study, we identified a few taxa associated with gavage treatments and weight-gain/-loss phenotypes, but unfortunately, we were unable to identify them to species or strain level. These taxa included Lachnospirales and Clostridia when mice on normal chow were gavaged with HFD-derived viromes (Fig. 5A and Fig. 5C), and Oscillospirales, Rhodospirales, Lachnospirales, and Peptostreptococcales when mice on HFD were gavaged with normal chow-derived viromes (Fig. 5B and Fig. 5D). While we were unable to identify these microbes beyond taxonomic order, other studies have found that microbes in these taxa were associated with weight gain/loss, suggesting that they could potentially be involved in our observed mouse phenotypes [46,47].

Our approach to study how virome gavage shapes the gut microbiota and affect lean/obese phenotypes in mice had several limitations. Firstly, because we used 16S rRNA amplicon sequencing, we were unable to make species-level classifications in most cases. In the future, we would prefer to use shotgun metagenomics in order to identify the microbes that were responsible for phenotype shifts at the species or strain level.

Another limitation was our decision to use whole viromes instead of individual viruses. While we were particularly interested in the collective effect of the virome on the bacterial biota and metabolism, this choice precluded our ability to identify specific viruses. We sequenced fecal viromes of recipient mice at several timepoints throughout the study, however we were unable to determine the virus groups that were responsible for the observed phenotypes (Fig. S6). We also sequenced the fecal and PEG-derived viromes from donor mice at the start of the experiment. While there are distinct differences between donor viromes, it is unclear whether these viruses were able to establish in recipient mice (Fig. S7). We believe it will take a dedicated study to confidently identify the viruses and bacterial strains that are responsible for the reported phenotypic changes.

As in any longitudinal study, the choice of timepoints to sample can limit findings. Our data suggest that the effect of virome perturbation on the bacterial biota manifested early in the study. Instead, a protocol that characterizes early timepoints more frequently may have yielded additional insights. Moreover, the purification protocol that we used to prepare virome gavages may have resulted in the loss of some viruses during PEG precipitation [48] which may have impacted the observed phenotypes. It is possible that fecal metabolites could have been carried over into the viromes, which could have affected the observed lean and obese phenotypes. However, we believe this is unlikely, as the PEG precipitation protocol involved centrifugation and resuspension in PBS to reduce metabolite carryover. Each of these purified viromes were subjected to culture to identify whether any cultivable bacteria were present, which provides broader confidence that these were pure viromes as we have used in prior studies [35].

The gut microbiome is a diverse and complicated ecological community with numerous types of microbes present. While much of the focus has been placed on the bacteria of the gut, which are known to shape a host metabolism, disease, and even behavior [49-51], it is becoming clear that the viral community also plays a role in many of these conditions [11,13,14]. Prior studies inform us that viruses in the gut have the capacity to shape their concomitant bacterial communities [22,25,35]. By extension, if bacterial communities can affect metabolic phenotypes [8], viromes can help to shape them as well. Our findings that gut-derived virome gavages are sufficient to alter the gut microbiota in mice and affect weight gain and loss contributes to previous demonstrations of the importance of the microbiome in shaping these phenotypes. As such, attention should be given to the potential effects of viruses in future microbiome transplant studies.

## Conclusions

In this study, we administered fecal virome transplants to mice on normal chow and high-fat diets in order to investigate whether viruses can affect the gut bacterial biota and elicit changes in host phenotypes. We find that virome gavages significantly altered the gut bacterial community and also affected mouse weight loss/gain; compared to counterparts on the same diet, mice receiving chow-derived viromes gained less weight and mice receiving high-fat diet-derived viromes gained more weight. Altogether, our work demonstrates that the gut virome is an important factor in shaping the gut bacterial biota and affecting host physiology.

## Abbreviations

HFD: high fat diet
Phage: bacteriophage
FVT: fecal virome transplant

## Declarations

### Animal Welfare Approval

All animal studies and research presented here were reviewed and approved by the Institutional Animal Care and Use Committee of the University of California, San Diego.

### Availability of Data and Material

All sequences included in this study have been deposited in the NCBI Sequence Read Archive under BioProject accession number PRJNA930016.

### Author Contributions

Conceived and designed project: JMB, DTP, BS, and CG.

Performed experiments: JMB, RL, YW, JC, PK, and LH.

Analyzed the data: JMB, RL, YW, DTP, BS, and XT.

Statistical analysis: JMB, TW, and XMT.

Wrote and edited the manuscript: JMB, RL, DTP, BS, JRM, CG, TW, and XMT.

Provided materials for the study: YW and BS.

All authors reviewed the manuscript.

## Acknowledgements

We thank Martin J. Blaser and Emma Allen Vercoe for valuable feedback on the conception and design of this project. We also thank Jonathan B. Shurin for contributing valuable input on analyses and Michael Overton for assistance with the Triton Shared Computing Cluster.

## References

1. Lloyd-Price, J., Abu-Ali, G., & Huttenhower, C. (2016). The healthy human microbiome. Genome Medicine, 8(1), 51. https://doi.org/10.1186/s13073-016-0307-y.

2. Shreiner, A. B., Kao, J. Y., & Young, V. B. (2015). The gut microbiome in health and in disease. Current Opinion in Gastroenterology, 31(1), 69–75. https://doi.org/10.1097/MOG.0000000000000139.

3. Gensollen, T., Iyer, S. S., Kasper, D. L., & Blumberg, R. S. (2016). How colonization by microbiota in early life shapes the immune system. Science, 352(6285), 539–544. https://doi.org/10.1126/science.aad9378.

4. Thaiss, C. A., Zmora, N., Levy, M., & Elinav, E. (2016). The microbiome and innate immunity. Nature, 535(7610), Article 7610. https://doi.org/10.1038/nature18847.

5. Rooks, M. G., & Garrett, W. S. (2016). Gut microbiota, metabolites and host immunity. Nature Reviews Immunology, 16(6), Article 6. https://doi.org/10.1038/nri.2016.42.

6. Carlson, A. L., Xia, K., Azcarate-Peril, M. A., Goldman, B. D., Ahn, M., Styner, M. A., Thompson, A. L., Geng, X., Gilmore, J. H., & Knickmeyer, R. C. (2018). Infant Gut Microbiome Associated With Cognitive Development. Biological Psychiatry, 83(2), 148–159. https://doi.org/10.1016/j.biopsych.2017.06.021.

7. Foster, J. A., & Neufeld, K. A. M. (2013). Gut–brain axis: How the microbiome influences anxiety and depression. Trends in Neurosciences, 36(5), 305–312. https://doi.org/10.1016/j.tins.2013.01.005.

8. Turnbaugh, P. J., Ley, R. E., Mahowald, M. A., Magrini, V., Mardis, E. R., & Gordon, J. I. (2006). An obesity-associated gut microbiome with increased capacity for energy harvest. Nature, 444(7122), 1027–1031. https://doi.org/10.1038/nature05414.

9. Lynch, S. V., & Pedersen, O. (2016). The Human Intestinal Microbiome in Health and Disease. New England Journal of Medicine, 375(24), 2369–2379. https://doi.org/10.1056/NEJMra1600266.

10. Lepage, P., Leclerc, M. C., Joossens, M., Mondot, S., Blottière, H. M., Raes, J., Ehrlich, D., & Doré, J. (2013). A metagenomic insight into our gut’s microbiome. Gut, 62(1), 146–158. https://doi.org/10.1136/gutjnl-2011-301805.

11. Carding, S. R., Davis, N., & Hoyles, L. (2017). Review article: The human intestinal virome in health and disease. Alimentary Pharmacology and Therapeutics, 46(9), 800–815. https://doi.org/10.1111/apt.14280.

12. Dahlman, S., Avellaneda-Franco, L., & Barr, J. J. (2021). Phages to shape the gut microbiota? Current Opinion in Biotechnology, 68, 89–95. https://doi.org/10.1016/j.copbio.2020.09.016.

13. Federici, S., Nobs, S. P., & Elinav, E. (2020). Phages and their potential to modulate the microbiome and immunity. Cellular and Molecular Immunology, August, 1–16. https://doi.org/10.1038/s41423-020-00532-4.

14. Mayneris-Perxachs, J., Castells-Nobau, A., Arnoriaga-Rodríguez, M., Garre-Olmo, J., Puig, J., Ramos, R., Martínez-Hernández, F., Burokas, A., Coll, C., Moreno-Navarrete, J. M., Zapata-Tona, C., Pedraza, S., Pérez-Brocal, V., Ramió-Torrentà, L., Ricart, W., Moya, A., Martínez-García, M., Maldonado, R., & Fernández-Real, J.-M. (2022). Caudovirales bacteriophages are associated with improved executive function and memory in flies, mice, and humans. Cell Host & Microbe, 30(3), 340–356.e8. https://doi.org/10.1016/j.chom.2022.01.013.

15. Khan Mirzaei, M., Khan, Md. A. A., Ghosh, P., Taranu, Z. E., Taguer, M., Ru, J., Chowdhury, R., Kabir, Md. M., Deng, L., Mondal, D., & Maurice, C. F. (2020). Bacteriophages Isolated from Stunted Children Can Regulate Gut Bacterial Communities in an Age-Specific Manner. Cell Host & Microbe, 27(2), 199-212.e5. https://doi.org/10.1016/j.chom.2020.01.004.

16. Minot, S., Sinha, R., Chen, J., Li, H., Keilbaugh, S. A., Wu, G. D., Lewis, J. D., & Bushman, F. D. (2011). The human gut virome: Inter-individual variation and dynamic response to diet. Genome Research, 21(10), 1616–1625. https://doi.org/10.1101/gr.122705.111.

17. Kim, M.S., & Bae, J.W. (2016). Spatial disturbances in altered mucosal and luminal gut viromes of diet-induced obese mice. Environmental Microbiology, 18(5), 1498–1510. https://doi.org/10.1111/1462-2920.13182.

18. Schulfer, A., Santiago-Rodriguez, T. M., Ly, M., Borin, J. M., Chopyk, J., Blaser, M. J., & Pride, D. T. (2020). Fecal Viral Community Responses to High-Fat Diet in Mice. MSphere, 5(1), e00833–19. https://doi.org/10.1128/mSphere.00833-19.

19. Quraishi, M. N., Widlak, M., Bhala, N., Moore, D., Price, M., Sharma, N., & Iqbal, T. H. (2017). Systematic review with meta-analysis: The efficacy of faecal microbiota transplantation for the treatment of recurrent and refractory Clostridium difficile infection. Alimentary Pharmacology & Therapeutics, 46(5), 479–493. https://doi.org/10.1111/apt.14201.

20. Federici, S., Kredo-Russo, S., Valdés-Mas, R., Kviatcovsky, D., Weinstock, E., Matiuhin, Y., Silberberg, Y., Atarashi, K., Furuichi, M., Oka, A., Liu, B., Fibelman, M., Weiner, I. N., Khabra, E., Cullin, N., Ben-Yishai, N., Inbar, D., Ben-David, H., Nicenboim, J., … Elinav, E. (2022). Targeted suppression of human IBD-associated gut microbiota commensals by phage consortia for treatment of intestinal inflammation. Cell, 185(16), 2879–2898.e24. https://doi.org/10.1016/j.cell.2022.07.003.

21. Duan, Y., Llorente, C., Lang, S., Brandl, K., Chu, H., Jiang, L., White, R. C., Clarke, T. H., Nguyen, K., Torralba, M., Shao, Y., Liu, J., Hernandez-Morales, A., Lessor, L., Rahman, I. R., Miyamoto, Y., Ly, M., Gao, B., Sun, W., … Schnabl, B. (2019). Bacteriophage targeting of gut bacterium attenuates alcoholic liver disease. Nature, 575(7783), Article 7783. https://doi.org/10.1038/s41586-019-1742-x.

22. Hsu, B. B., Gibson, T. E., Yeliseyev, V., Liu, Q., Lyon, L., Bry, L., Silver, P. A., & Gerber, G. K. (2019). Dynamic Modulation of the Gut Microbiota and Metabolome by Bacteriophages in a Mouse Model. Cell Host and Microbe, 25(6), 803–814.e5. https://doi.org/10.1016/j.chom.2019.05.001.

23. Draper, L. A., Ryan, F. J., Dalmasso, M., Casey, P. G., McCann, A., Velayudhan, V., Ross, R. P., & Hill, C. (2020). Autochthonous faecal viral transfer (FVT) impacts the murine microbiome after antibiotic perturbation. BMC Biology, 18(1), 173. https://doi.org/10.1186/s12915-020-00906-0.

24. Rasmussen, T. S., Mentzel, C. M. J., Kot, W., Castro-Mejía, J. L., Zuffa, S., Swann, J. R., Hansen, L. H., Vogensen, F. K., Hansen, A. K., & Nielsen, D. S. (2020). Faecal virome transplantation decreases symptoms of type 2 diabetes and obesity in a murine model. Gut, 69(12), 2122–2130. https://doi.org/10.1136/gutjnl-2019-320005

25. Mu, A., McDonald, D., Jarmusch, A. K., Martino, C., Brennan, C., Bryant, M., Humphrey, G. C., Toronczak, J., Schwartz, T., Nguyen, D., Ackermann, G., D’Onofrio, A., Strathdee, S. A., Schooley, R. T., Dorrestein, P. C., Knight, R., & Aslam, S. (2021). Assessment of the microbiome during bacteriophage therapy in combination with systemic antibiotics to treat a case of staphylococcal device infection. Microbiome, 9(1), 1–8. https://doi.org/10.1186/s40168-021-01026-9.

26. Lourenço, M., Chaffringeon, L., Lamy-Besnier, Q., Titécat, M., Pédron, T., Sismeiro, O., Legendre, R., Varet, H., Coppée, J.-Y., Bérard, M., De Sordi, L., & Debarbieux, L. (2022). The gut environment regulates bacterial gene expression which modulates susceptibility to bacteriophage infection. Cell Host & Microbe, 30(4), 556–569.e5. https://doi.org/10.1016/j.chom.2022.03.014

27. Hoyles, L., McCartney, A. L., Neve, H., Gibson, G. R., Sanderson, J. D., Heller, K. J., & van Sinderen, D. (2014). Characterization of virus-like particles associated with the human faecal and caecal microbiota. Research in Microbiology, 165(10), 803–812. https://doi.org/10.1016/j.resmic.2014.10.006.

28. Klindworth, A., Pruesse, E., Schweer, T., Peplies, J., Quast, C., Horn, M., & Glöckner, F. O. (2013). Evaluation of general 16S ribosomal RNA gene PCR primers for classical and next-generation sequencing-based diversity studies. Nucleic Acids Research, 41(1), e1. https://doi.org/10.1093/nar/gks808.

29. Bolyen, E., Rideout, J. R., Dillon, M. R., Bokulich, N. A., Abnet, C. C., Al-Ghalith, G. A., Alexander, H., Alm, E. J., Arumugam, M., Asnicar, F., Bai, Y., Bisanz, J. E., Bittinger, K., Brejnrod, A., Brislawn, C. J., Brown, C. T., Callahan, B. J., Caraballo-Rodríguez, A. M., Chase, J., … Caporaso, J. G. (2019). Author Correction: Reproducible, interactive, scalable and extensible microbiome data science using QIIME 2. Nature Biotechnology, 37(9), Article 9. https://doi.org/10.1038/s41587-019-0252-6.

30. Callahan, B. J., McMurdie, P. J., Rosen, M. J., Han, A. W., Johnson, A. J. A., & Holmes, S. P. (2016). DADA2: High-resolution sample inference from Illumina amplicon data. Nature Methods, 13(7), Article 7. https://doi.org/10.1038/nmeth.3869.

31. Quast, C., Pruesse, E., Yilmaz, P., Gerken, J., Schweer, T., Yarza, P., Peplies, J., & Glöckner, F. O. (2013). The SILVA ribosomal RNA gene database project: Improved data processing and web-based tools. Nucleic Acids Research, 41(D1), D590–D596. https://doi.org/10.1093/nar/gks1219.

32. Bisanz, J. E. (2018). Qiime2R; Importing QIIME2 artifacts and associated data into R sessions. Version 0.99. https://github.com/jbisanz/qiime2R.

33. Wickham, H. (2016). ggplot2: Elegant Graphics for Data Analysis. Springer-Verlag New York. ISBN 978-3-319-24277-4, https://ggplot2.tidyverse.org.

34. R Core Team (2021). R: A language and environment for statistical computing. R Foundation for Statistical Computing, Vienna, Austria. https://www.R-project.org/.

35. Attai, H., Wilde, J., Liu, R., Chopyk, J., Garcia, A. G., Allen-Vercoe, E., & Pride, D. (2022). Bacteriophage-Mediated Perturbation of Defined Bacterial Communities in an In Vitro Model of the Human Gut. Microbiology Spectrum, 10(3), e01135–22. https://doi.org/10.1128/spectrum.01135-22.

36. Conceição-Neto, N., Matthijnssens, J., Zeller, M., Lefrère, H., De Bruyn, P., Beller, L., Deboutte, W., Yinda, C. K., Lavigne, R., Maes, P., & Van Ranst, M. (2016). NetoVIR: a reproducible protocol for virome analysis. Protocol Exchange 5:16532. https://doi.org/10.1038/protex.2016.029.

37. Santiago-Rodriguez, T. M., Ly, M., Daigneault, M. C., Brown, I. H. L., McDonald, J. A. K., Bonilla, N., Vercoe, E. A., & Pride, D. T. (2015). Chemostat culture systems support diverse bacteriophage communities from human feces. Microbiome, 3(1), 58. https://doi.org/10.1186/s40168-015-0124-3

38. Tang, W., He, H., & Tu, X. M. (2012). Applied categorical and count data analysis. CRC Press.

39. Oksanen, A. J. R. I, Blanchet, F. G., Friendly, M., Kindt, R., Legendre, P., McGlinn, D., Minchin, P. R., O’Hara, R. B., Simpson, G. L., Solymos, P., and Stevens, M. H. H. (2020). vegan: Community Ecology Package. R package version 2.5–7. https://github.com/vegandevs/vegan.

40. Halekoh U, Højsgaard S, Yan J (2006). The R Package geepack for Generalized Estimating Equations. Journal of Statistical Software, 15(2), 1–11.

41. Holm, S. (1979). A simple sequentially rejective multiple test procedure. Scandinavian journal of statistics, 65–70.

42. Martino, C., Morton, J. T., Marotz, C. A., Thompson, L. R., Tripathi, A., Knight, R., & Zengler, K. (2019). A Novel Sparse Compositional Technique Reveals Microbial Perturbations. MSystems, 4(1), e00016–19. https://doi.org/10.1128/mSystems.00016-19.

43. Ly, M., Abeles, S. R., Boehm, T. K., Robles-Sikisaka, R., Naidu, M., Santiago-Rodriguez, T., & Pride, D. T. (2014). Altered Oral Viral Ecology in Association with Periodontal Disease. MBio, 5(3), e01133–14. https://doi.org/10.1128/mBio.01133-14.

44. Norman, J. M., Handley, S. A., Baldridge, M. T., Droit, L., Liu, C. Y., Keller, B. C., Kambal, A., Monaco, C. L., Zhao, G., Fleshner, P., Stappenbeck, T. S., McGovern, D. P. B., Keshavarzian, A., Mutlu, E. A., Sauk, J., Gevers, D., Xavier, R. J., Wang, D., Parkes, M., & Virgin, H. W. (2015). Disease-Specific Alterations in the Enteric Virome in Inflammatory Bowel Disease. Cell, 160(3), 447–460. https://doi.org/10.1016/j.cell.2015.01.002.

45. Wang, L., Mazagova, M., Pan, C., Yang, S., Brandl, K., Liu, J., Reilly, S. M., Wang, Y., Miao, Z., Loomba, R., Lu, N., Guo, Q., Liu, J., Yu, R. T., Downes, M., Evans, R. M., Brenner, D. A., Saltiel, A. R., Beutler, B., & Schnabl, B. (2019). YIPF6 controls sorting of FGF21 into COPII vesicles and promotes obesity. Proceedings of the National Academy of Sciences of the United States of America, 116(30), 15184–15193. https://doi.org/10.1073/pnas.1904360116

46. Hossain, M. M., Begum, M., and Kim, I. H. (2015). Effect of *Bacillus subtilis*, *Clostridium butyricum*, and *Lactobacillus acidophilus* endospores on growth performance, nutrient digestibility, meat quality, relative organ weight, microbial shedding and excreta noxious gas emission in broilers. Veterinarni Medicina, 60(2), 77–86. https://doi.org/10.17221/7981-VETMED.

47. Vacca, M., Celano, G., Calabrese, F. M., Portincasa, P., Gobbetti, M., & De Angelis, M. (2020). The Controversial Role of Human Gut Lachnospiraceae. Microorganisms, 8(4), Article 4. https://doi.org/10.3390/microorganisms8040573.

48. Carroll-Portillo, A., Coffman, C. N., Varga, M. G., Alcock, J., Singh, S. B., & Lin, H. C. (2021). Standard Bacteriophage Purification Procedures Cause Loss in Numbers and Activity. Viruses, 13(2), Article 2. https://doi.org/10.3390/v13020328

49. Kamada, N., Chen, G. Y., Inohara, N., & Núñez, G. (2013). Control of pathogens and pathobionts by the gut microbiota. Nature Immunology, 14(7), Article 7. https://doi.org/10.1038/ni.2608

50. Gilbert, J. A., Blaser, M. J., Caporaso, J. G., Jansson, J. K., Lynch, S. V., & Knight, R. (2018). Current understanding of the human microbiome. Nature Medicine, 24(4), Article 4. https://doi.org/10.1038/nm.4517

51. Fan, Y., & Pedersen, O. (2021). Gut microbiota in human metabolic health and disease. Nature Reviews Microbiology, 19(1), Article 1. https://doi.org/10.1038/s41579-020-0433-9

