## Supplemental Material for "Fecal virome transplantation is sufficient to alter fecal microbiota and drive lean and obese body phenotypes in mice"

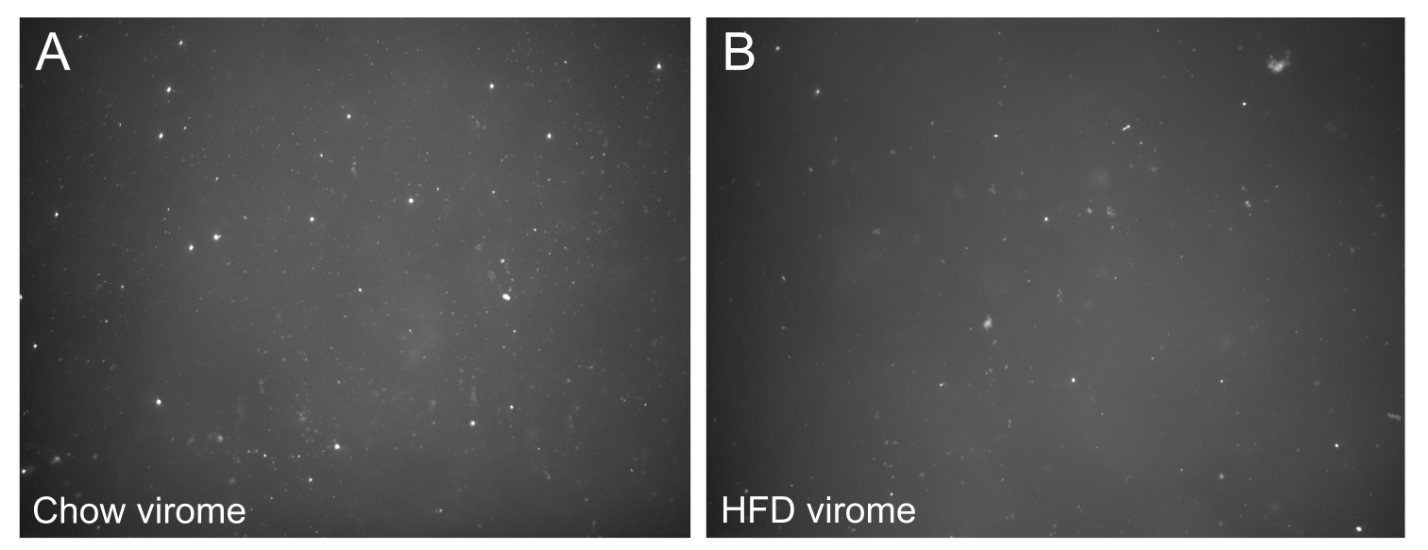

**Figure S1.** Representative epifluorescence microscopy images of PEG-precipitated chow (Panel A) and HFD (Panel B) viromes used for FVT gavage. Samples were imaged at 1000× and manually enumerated in 3–5 fields of view per slide. The concentration of VLPs in each sample was calculated by N_v_ = P_t_ / F_t_ * A_t_ / A_f_ / V_t_ where N_v_ is VLPs per mL, P_t_ is the number of VLPs counted, F_t_ is the number of fields of view counted, A_t_ is the total area of the Anodisc, A_f_ is the area of each field of view, and V_t_ is the volume of sample that was filtered onto the Anodisc. No significant difference was found between chow and HFD virome densities.

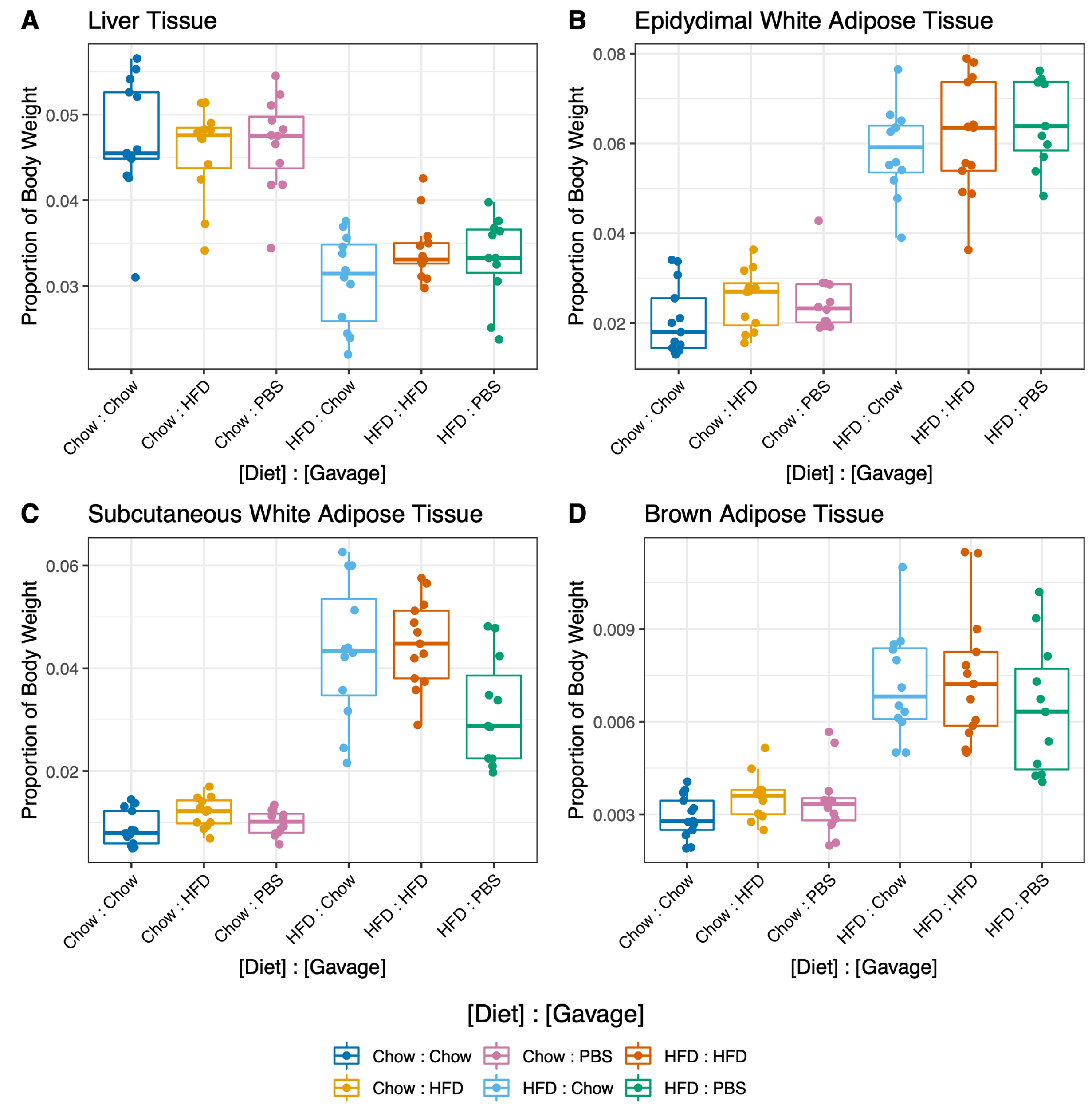

**Figure S2.** Various tissue weights relative to body mass. Tissues were harvested from individual mice after 4 weeks of gavage treatment. Tissues included the liver (A), epidydimal white adipose tissue (B), subcutaneous white adipose tissue (C), and brown adipose tissue (D). Diet had a significant effect on all tissue weights (p<2.2e-16) and the effect of gavage was not significant (GEE GLM Tissue proportion ~ Diet + Gavage). Data from PBS controls were omitted from statistical models.

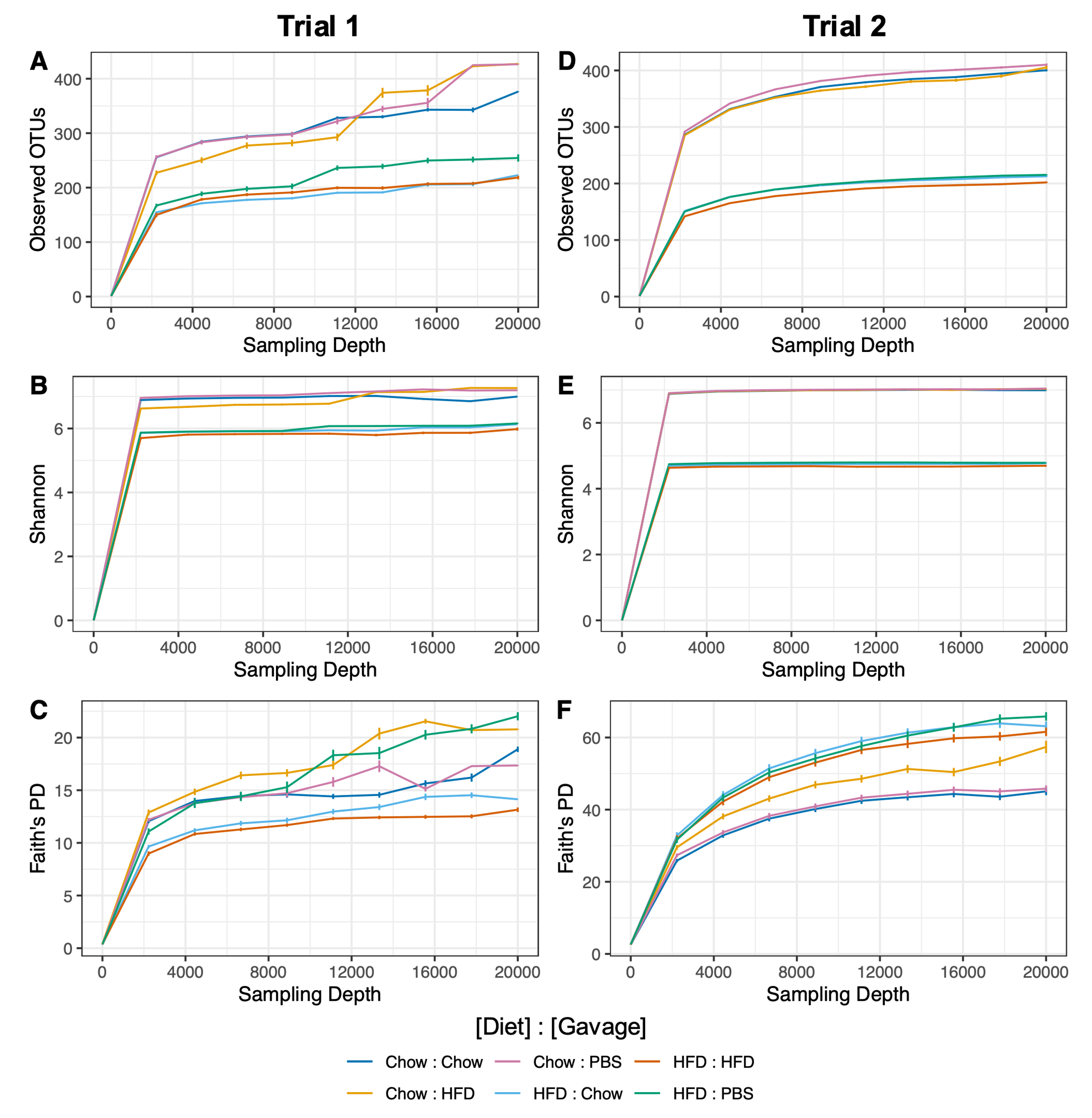

**Figure S3.** Rarefaction curves for all samples grouped by trial and alpha diversity metric. Trial 1 (left) shows curves for Observed OTUs, Shannon, and Faith’s phylogenetic distance (Panels A, B, and C, respectively). Trial 2 (right) shows curves for Observed OTUs, Shannon, and Faith’s phylogenetic distance (Panels D, E, and F, respectively). Lines represent means and error bars indicate standard error. Colors represent unique treatments (pair of diet and gavage) indicated in the legend.

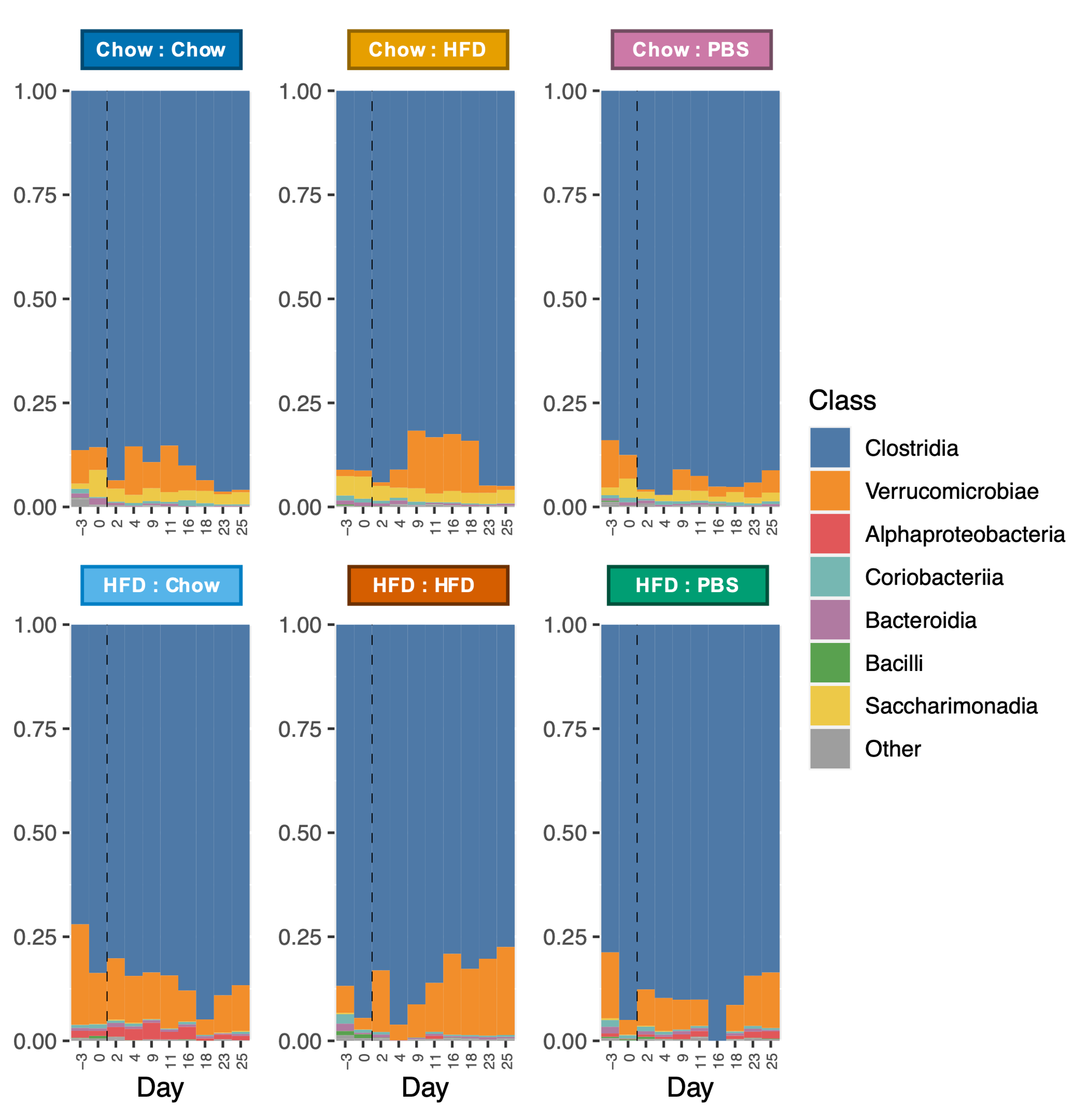

**Figure S4.** Taxonomy stacked bar plots indicating proportions of bacterial classes throughout the trial 1 experiment. Each panel represents a cage of 4 mice. Labels above panels indicate treatment group (Diet : Gavage). The dashed vertical line indicates start of gavage treatments. Treatment mice were gavaged with viromes from a separate group of donor mice on a chow diet or high-fat diet, or with a PBS control gavage (n=6 cages, one per treatment).

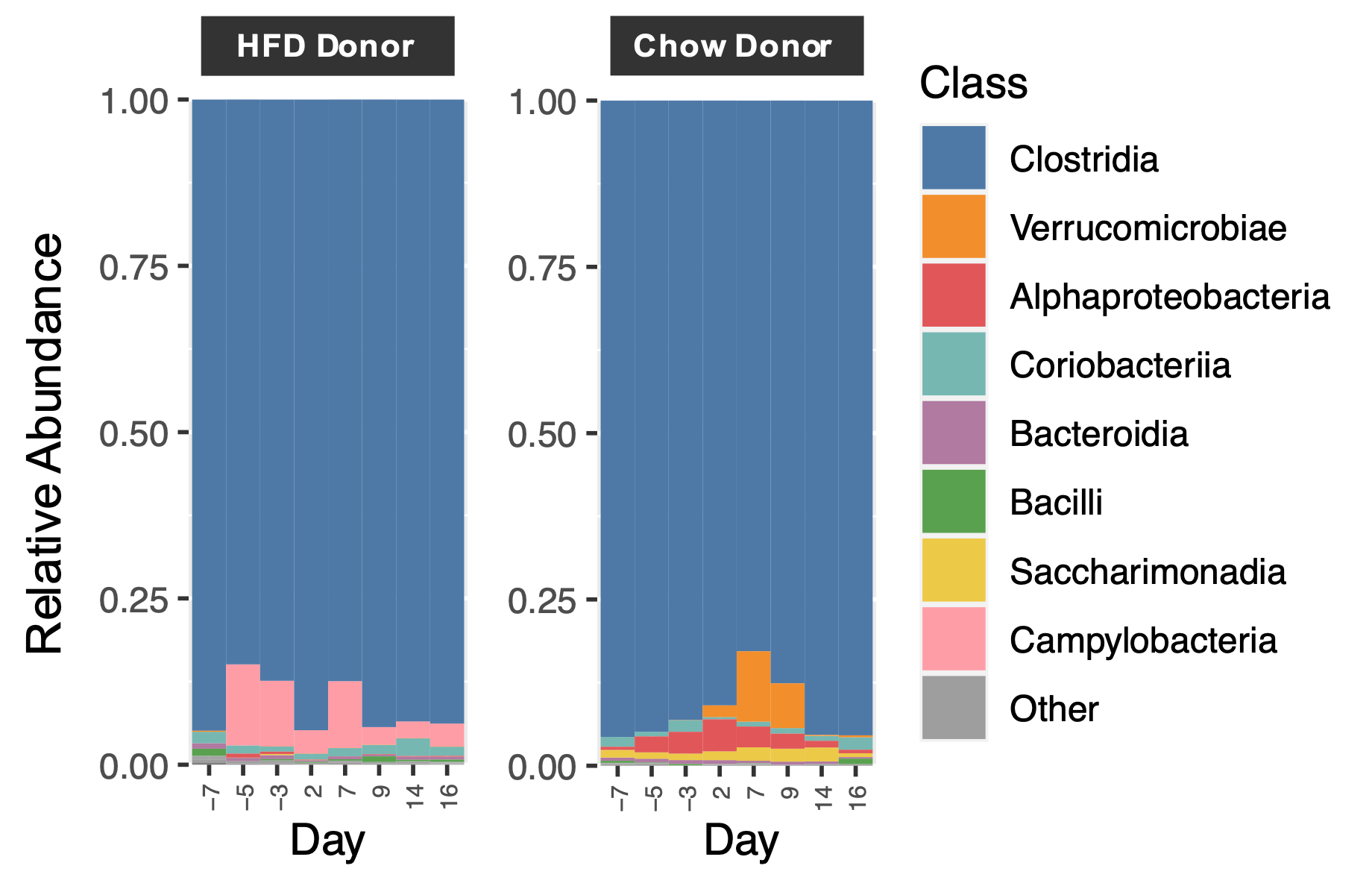

**Figure S5.** Taxonomy stacked bar plots indicating proportions of bacterial classes in donor mice. Each panel represents a cage of 4 mice. Labels above panels indicate donor diet. Viromes were prepared from donors and administered to respective treatment mice one week later (e.g., donor viromes from day -7 were administered to recipient mice on day 0).

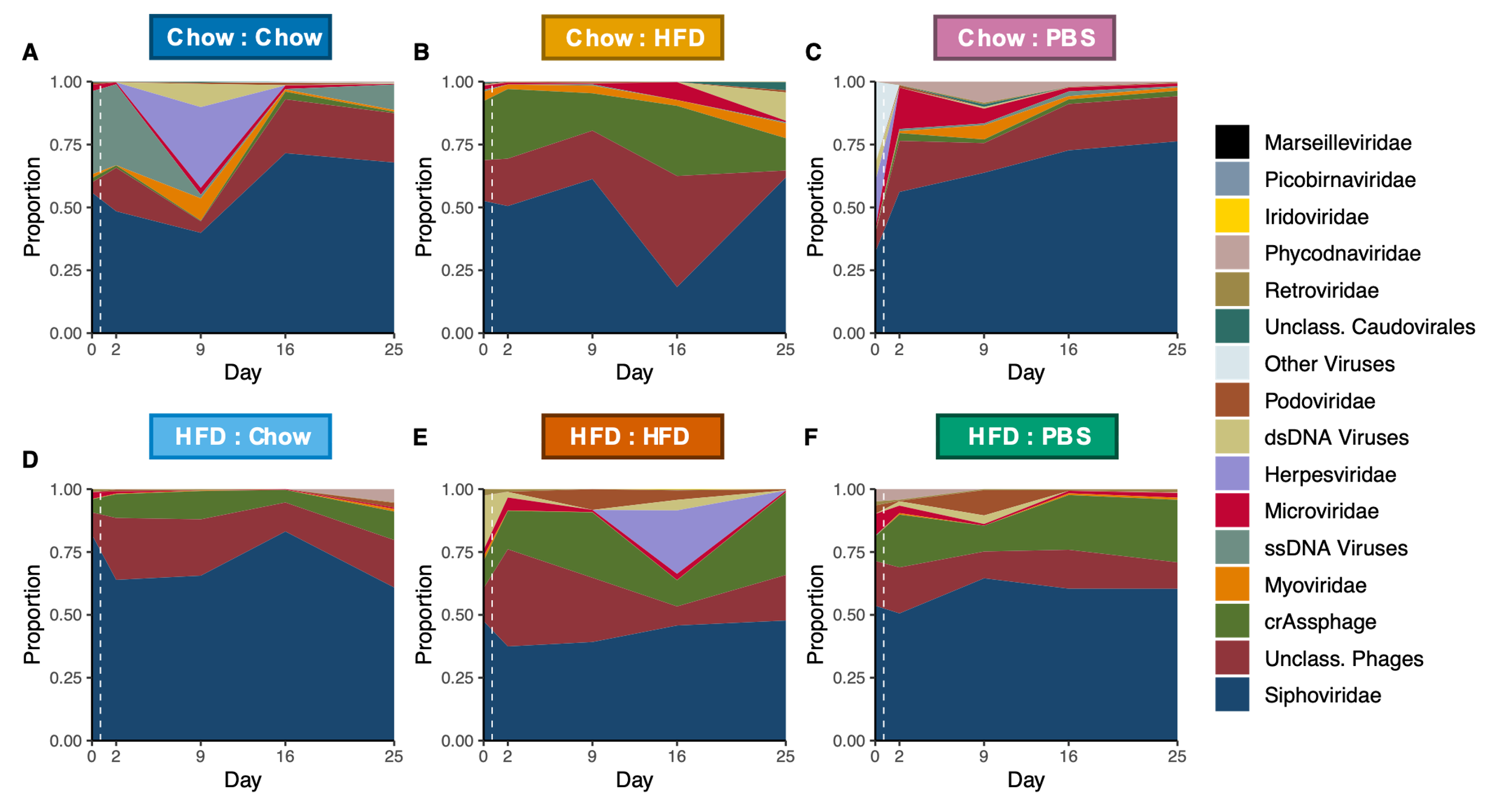

**Figure S6.** Taxonomy stacked area plots indicating proportions of virus groups sampled on days 0, 2, 9, 16, and 25 of the trial 2 experiment. Each panel represents averages sampled from 3 replicate cages. Labels above panels indicate treatment group (Diet : Gavage). The dashed vertical line indicates the start of gavage treatments. Treatment mice were gavaged with viromes from a separate group of donor mice on a chow diet or high-fat diet (see Fig. S7), or with a PBS control gavage (n=18 cages, 3 per treatment).

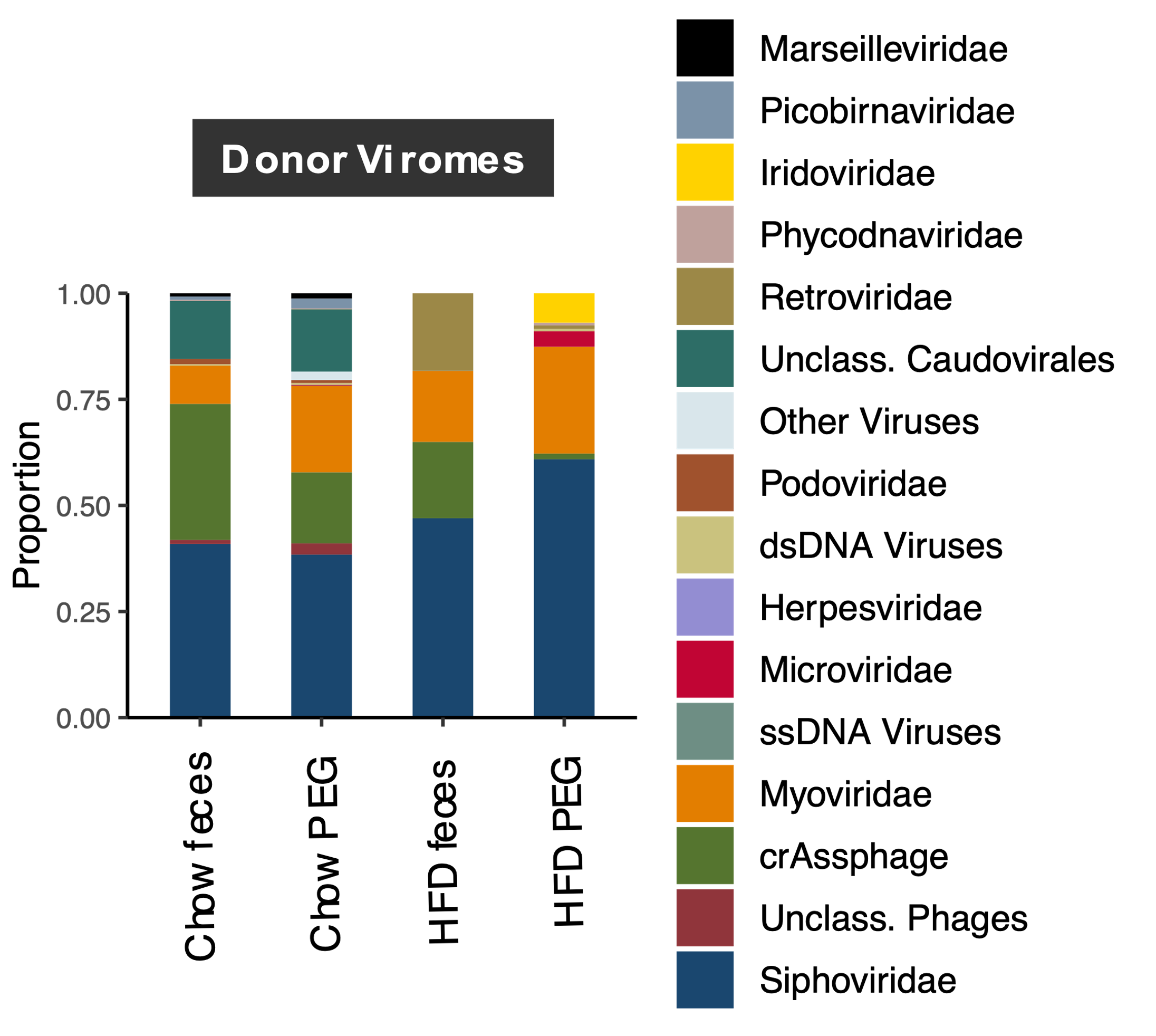

**Figure S7.** Taxonomy stacked bar plots indicating proportions of virus groups sampled from donor mice on day 0. Viromes were extracted, sequenced, and analyzed from fecal samples (as performed for treatment mouse samples, see Fig. S6), as well as from PEG-precipitated samples that were derived from these fecal samples and which were used to gavage treatment mice.
